# Downregulation of O-GlcNAcylation enhances etoposide-induced p53-mediated apoptosis in HepG2 human liver cancer cells

**DOI:** 10.1101/2024.11.29.626030

**Authors:** Jaehoon Lee, Gi-Bang Koo, Jihye Park, Byung-Cheol Han, Mijin Kwon, Seung-Ho Lee

## Abstract

Etoposide, an anticancer drug that inhibits topoisomerase II, is commonly used in combination chemotherapy. However, the impact of O-GlcNAcylation regulation on etoposide’s anticancer effects has rarely been investigated. This study evaluated the effect of etoposide on cellular O-GlcNAcylation and whether modulating this process enhances etoposide-induced apoptosis. O-GlcNAc expression was measured after 24 h of etoposide treatment, and the effect of O-GlcNAc transferase (OGT) inhibition by OSMI-1 on etoposide’s anticancer activity in HepG2 human liver cancer cells was quantitatively analyzed. Additionally, molecular analyses were used to confirm that the observed effects were mediated by p53-induced apoptosis. Etoposide reduced O-GlcNAcylation in a dose-dependent manner without directly interacting with OGT. Cotreatment with 20 μM of OSMI-1 lowered the IC_50_ value for cell viability by approximately 1.64-fold to 60.68 μM and increased the EC_50_ value for cytotoxicity by around 4.07-fold to 43.41 μM. Furthermore, this synergistic effect was linked to the activation of the p53/caspase-3/PARP1 pathway. These findings suggest that downregulating O-GlcNAcylation may enhance the efficacy of etoposide-based chemotherapy and help overcome tumor resistance.

**Graphical Abstract:** Etoposide, a topoisomerase II inhibitor, reduces O-GlcNAcylation in HepG2 liver cancer cells. Further inhibition of O-GlcNAc transferase by OSMI-1 enhanced etoposide-induced apoptosis, lowering the IC50 for viability and increasing the EC50 for cytotoxicity. This synergy was mediated by the p53, caspase-3/7, PARP1 pathway, suggesting that targeting O-GlcNAcylation may improve etoposide efficacy.

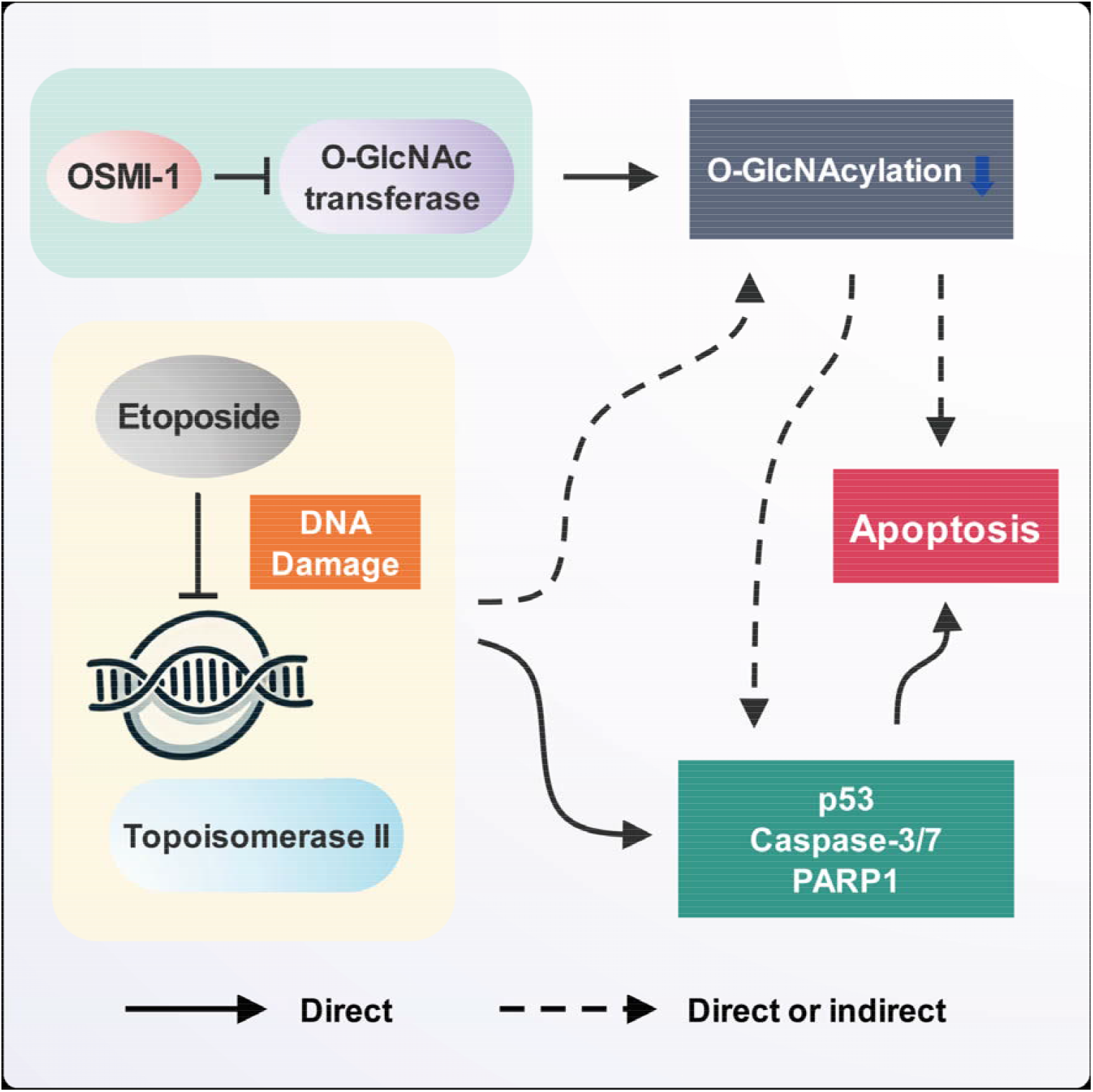

## Introduction

O-linked β-N-acetylglucosamine modification (O-GlcNAcylation) is a well-known post-translational modification that occurs in response to endogenous or exogenous factors, primarily in the nucleus, cytoplasm, and mitochondria [1]. O-GlcNAcylation plays a critical role in regulating transcription, metabolism, cell signaling, and physiological activity [2–4]. Prior studies have linked precise O-GlcNAcylation regulation to immune [5, 6] and cardiac functions [7], metabolic diseases [8], autoimmune disorders [9], and neurodegenerative diseases, such as Parkinson’s and Alzheimer’s disease [10]. Interestingly, O-GlcNAcylation is regulated by two highly conserved enzyme pairs: O-GlcNAc transferase (OGT) and protein O-GlcNAcase (OGA) [11]. Therefore, OGT and OGA are targets for disease treatment studies.

OGT function and O-GlcNAcylation regulation are involved in tumor formation, survival, metastasis, and angiogenesis. OGT activity and O-GlcNAcylation are increased in most malignant tumors and correlate positively with tumor progression and recurrence [12]. For example, studies have shown that OGT inhibition reduces cell cycle progression in a c-MYC–dependent manner in prostate cancer cells [13] and decreases epithelial–mesenchymal transition, metastasis, and invasion *in vivo* [14, 15]. Therefore, modulating O-GlcNAcylation by inhibiting OGT activity may be a promising target for cancer treatment.

Etoposide, an anticancer drug, forms a ternary complex with topoisomerase II (Topo II) and DNA, inducing apoptosis by inhibiting DNA strand religation [16]. This mechanism makes etoposide effective against various cancers, particularly those with rapid cell division. p53, a key transcription factor in apoptosis, is mutated in over 50% of human tumors [17]. When p53 is dysfunctional, cell cycle regulation and apoptosis are impaired, leading to uncontrolled cell proliferation and tumor development [18]. Thus, p53 is a highly effective target for cancer therapy. Chemotherapeutic drugs, such as etoposide and cisplatin, increase p53 expression, which in turn regulates the expression of *CDKN1A, GADD45A, BAX*, and other genes to induce cancer cell death [19–21]. However, additional resistance or the tumor microenvironment can allow cancer cells to evade the p53-mediated apoptosis pathway. For example, in HepG2 liver cancer cells, hypoxia conditions silence etoposide-induced p53 expression [22–24], suggesting that etoposide alone may have limited effectiveness *in vivo*.

Previous studies have shown that OGT inhibition with OSMI-1, an OGT inhibitor, increases sensitivity to the anticancer drug doxorubicin in HepG2 human liver cancer cells and PC-3 prostate cancer cells [25, 26]. Additionally, downregulating OGT activity using siOGT has been found to enhance the apoptotic response to bortezomib in MCF-7 and T47D human breast cancer cells [27]. However, the effect of O-GlcNAcylation regulation on etoposide’s anticancer activity remains unexplored.

In this study, we investigated how O-linked β-N-acetylglucosamine (O-GlcNAc) expression is regulated by etoposide treatment and demonstrated that further downregulation of O-GlcNAcylation enhances the anticancer effect of etoposide in multiple human cancer cells. We also confirmed that these effects occur through the p53-mediated apoptosis pathway. Our findings suggest that reducing O-GlcNAcylation could help overcome tumor resistance and improve therapeutic outcomes in etoposide-based chemotherapy.

## Methods

### General materials

Etoposide and Thiamet G were obtained from MedChemExpress (Monmouth Junction, NJ, USA). OSMI-1 and phalloidin-iFluor™ 555 conjugate were purchased from GlpBio (Montclair, CA, USA) and Cayman Chemical (Ann Arbor, MI, USA), respectively. Crystal violet solution, Tween 20, and bisBenzimide H 33342 trihydrochloride (Hoechst 33342) were acquired from Sigma-Aldrich (Saint Louis, MO, USA). Antibodies were sourced from the following companies: O-GlcNAc (monoclonal; #82332) for western blot analysis from Cell Signaling Technology (Danvers, MA, USA); O-GlcNAc (RL2; monoclonal; #MA1-072) antibodies for immunocytochemistry from Thermo Fisher Scientific (Waltham, MA, USA); OGT (ARC0790; monoclonal; #A3501), OGA (ARC65434; monoclonal; #A24124), PARP1 (ARC0075; monoclonal; #A19596), and cleaved-PARP1 (ARC0091; monoclonal; #A19612) antibodies from ABclonal Technology (Woburn, MA, USA); and β-actin (polyclonal; #ab8227) and p53 (PAb421; monoclonal; #NBP2-62555) antibodies from Abcam (Cambridge, United Kingdom) and Novus Biologicals (Centennial, CO, USA), respectively.

### Cell culture

HepG2 and Hep3B human liver hepatocellular carcinoma, A549 human lung carcinoma, and Caco-2 human colorectal adenocarcinoma cell lines were obtained from the American Type Culture Collection (ATCC; Manassas, VA, USA). HepG2, Hep3B, Caco-2 cells were cultured in Eagle’s Minimum Essential Medium (EMEM; ATCC, Chicago, IL, USA) containing 1.0 g/L D-glucose, 10% fetal bovine serum (FBS), 1% penicillin/streptomycin (P/S), 1.5 g/L sodium bicarbonate, 292.0 mg/L L-glutamine, and 110.0 mg/L sodium pyruvate. A549 cells were cultured in Kaighn’s Modification of Ham’s F-12 Medium (ATCC) containing 1.26 g/L D-glucose, 10% FBS, 1% P/S, 1.5 g/L sodium bicarbonate, 292.2 mg/L L-glutamine, and 220.0 mg/L sodium pyruvate. All cells were cultured at 37°C in an incubator with a 5% CO_2_ humidified atmosphere.

#### Cell viability assay

HepG2 cells were seeded in 96-well plates at 1.0 × 10^4^ cells per well and maintained for 24 h. The medium was replaced with different concentrations of drugs, and after 24 h, cell viability was assessed using the D-Plus™ CCK Cell Viability Assay Kit (Dongin LS, Seoul, South Korea).

### Crystal violet staining

To evaluate the effects of etoposide and OSMI-1 on HepG2 growth, cells were seeded in EMEM in 48-well plates at 5.0 × 10^4^ cells per well. After 24 h, the medium was replaced with the drugs. Following 24 h of treatment, the cells were stained with crystal violet for 20 min and then washed thoroughly with phosphate-buffered saline (PBS). ImageJ software (version 1.54g) was used to quantify crystal violet intensity.

### Western blot analysis

HepG2 cells were harvested and lysed on ice for 5 min using Cell Culture Lysis 1X Reagent (Promega, Madison, WI, USA) containing a mixture of protease and phosphatase inhibitors (MedChemExpress, Monmouth Junction, NJ, USA). The insoluble debris was removed via centrifugation at 15,000 × *g* for 20 min at 4°C, and the total protein concentration of the isolated supernatants was quantified using the Pierce^TM^ BCA Protein Assay Kit (Thermo Fisher Scientific). Proteins (10□20 μg) were separated using the Bolt™ Bis-Tris Plus Gel system (Thermo Fisher Scientific) and transferred to polyvinylidene difluoride membranes (Bio-Rad, Hercules, CA, USA). The membranes were incubated in 0.05% Tween 20 Tris-buffered saline (TBST) containing 3%–5% bovine serum albumin (BSA) for 1 h at room temperature, followed by incubation overnight at 4°C in 5% BSA–TBST with diluted primary antibodies. The membranes were then washed three times with TBST and incubated with horseradish peroxidase–conjugated secondary antibodies in 5% BSA–TBST for 1 h at room temperature. Protein signals were developed on the membranes using ECL substrates (Thermo Fisher Scientific) and analyzed using Fusion FX6.0 (Vilber, Collégien, France).

### Immunocytochemistry

For immunocytochemistry, HepG2 cells were cultured with drugs at an appropriate density on gelatin-coated coverslips (SPL Life Sciences, Pocheon-si, South Korea). Cells were fixed with 4% paraformaldehyde for 20□min and permeabilized with 0.2% Triton X-100 for 5 min. After blocking with 10% normal donkey serum (Abcam) for 1□h, the coverslips were incubated overnight with primary antibodies diluted in 10% normal donkey serum. Subsequently, the coverslips were washed three times with PBS containing 0.1% Tween 20 (PBST) and incubated with Alexa Fluor 488–conjugated secondary antibodies for 30□min. After washing with PBST, the coverslips were incubated with phalloidin-iFluor™ 555 conjugate and Hoechst 33342 for 10 min, followed by thorough washing. Finally, the coverslips were mounted on slides using ProLong™ Glass Antifade Mountant (Thermo Fisher Scientific), and the slides were analyzed using LSM 900 with Airyscan 2 confocal microscopes (Carl Zeiss, Oberkochen, Germany).

### OGT activity assay

OGT activity was assessed using the Uridine diphosphate (UDP)-Glo™ Glycosyltransferase Assay Kit (Promega) according to the manufacturer’s protocol. UDP-GlcNAc (Promega) was utilized as the UDP–sugar substrate, while the OGT peptide substrate (KKKYPGGSTPVSSANMM) was sourced from AnaSpec (Fremont, CA, USA). Recombinant human OGT protein was obtained from R&D Systems (Minneapolis, MN, USA). The enzymatic reactions were performed in 1X OGT reaction buffer, consisting of 25 mM Tris (pH 7.5), 12.5 mM MgCl□, 0.06 mg/mL BSA, and 1 mM dithiothreitol.

### Fluorescence-activated cell sorting

HepG2 cells were seeded at 1.0 × 10^6^ cells per well in 6-well plates in the medium described above. After 24 h, the medium was replaced with fresh medium containing the desired concentrations of etoposide and OSMI-1, after which it was maintained for 24 h. Cells were then harvested and stained using the Annexin V Apoptosis Detection Kit with 7-AAD (STEMCELL Technologies, Vancouver, BC, Canada), following the manufacturer’s instructions, and analyzed using the FACSLyric™ System (BD Biosciences, Franklin Lakes, NJ, USA).

### Annexin V binding and membrane integrity assay

To assess the effect of drugs on apoptosis and necrosis in HepG2 cells, the RealTime-Glo™ Annexin V Apoptosis and Necrosis Assay Kit (Promega) was used. Briefly, HepG2 cells were seeded in solid white 96-well plates at 1.0 × 10^4^ cells per well and maintained for 24 h. The medium was replaced with fresh medium containing the drug and kit components according to the manufacturer’s instructions. Luminescence and fluorescence signals were measured using the GloMax® Discover Microplate Reader (Promega).

### TUNEL assay

HepG2 cells cultured in 6-well plates were fixed with 4% formaldehyde for 30 min and permeabilized with 0.2% Triton X-100 and 0.5% BSA in PBS. The cells were then incubated with TUNEL reaction mixture (Abbkine, Atlanta, GA, USA) for 2 h at 37°C, followed by phalloidin-iFluor™ 555 conjugate and 4’, 6-diamidino-2-phenylindole (i.e., DAPI) for an additional 10 min at room temperature. After thorough washing with PBS, the cells were analyzed using LSM 900 with Airyscan 2 confocal microscopes.

### Viability (GF-AFC), cytotoxicity (bis-AAF-R110), and caspase-3/7 activity assay

To evaluate the effects of drugs on viability, cytotoxicity, and caspase-3/7 activity, HepG2 cells were assessed using the ApoTox-Glo™ Triplex Assay Kit (Promega). Briefly, HepG2 cells were seeded in solid white 96-well plates at 1.0 × 10^4^ cells per well and maintained for 24 h. The medium was replaced with different drug concentrations, and after 24 h, GF-AFC substrate and bis-AAF-R110 substrate were added according to the manufacturer’s protocol. After 2 h, fluorescence analysis (viability: 400/505 nm excitation/emission; cytotoxicity: 485/520 nm excitation/emission) was conducted. A luminogenic caspase-3/7 substrate was then added, and the plates were incubated for an additional 2 h at 37°C. Luminescence and fluorescence signals were measured using the GloMax® Discover Microplate Reader (Promega).

### Statistical analysis

Statistical analysis was performed using SPSS Statistics version 20.0 (IBM Corp., Armonk, NY, USA) and GraphPad Prism 9 (GraphPad Software Inc., La Jolla, CA, USA). Data are presented as means ± standard deviations (SDs). Analysis of variance (ANOVA) with the post-hoc Tukey HSD test was used for data analysis. *P* < 0.05 was considered statistically significant.

## Results

### OGT inhibitor enhances sensitivity to etoposide in HepG2 cells

The effect of etoposide on the viability of HepG2 cells in the presence of the OGT inhibitor OSMI-1 was assessed using the CCK cell viability assay 24 h after treatment. Results showed that coadministration of etoposide with 20 μM OSMI-1 significantly reduced cell viability at all concentrations compared with etoposide alone (Fig. 1A). These findings were further validated through crystal violet staining (Fig. 1B and C). Viability analysis using GF-AFC showed that cotreatment with 20 μM OSMI-1 reduced the IC_50_ value of etoposide in HepG2 cells by approximately 1.64-fold, from 99.59 to 60.68 μM (Fig. 1D). Additionally, the membrane integrity assay, which serves as a cytotoxicity index, revealed that the EC_50_ value of etoposide decreased approximately 4.07-fold from 43.41 to 176.63 μM, following cotreatment with 20 μM OSMI-1 (Fig. 1E). The same trend was observed in other human cancer cell lines, including Hep3B, A549, and Caco-2, where cotreatment with etoposide and OSMI-1 resulted in a more significant reduction in cell viability (Fig. 1F). These results suggest that the anticancer effects of etoposide are influenced by O-GlcNAc regulation.

**Fig. 1.**
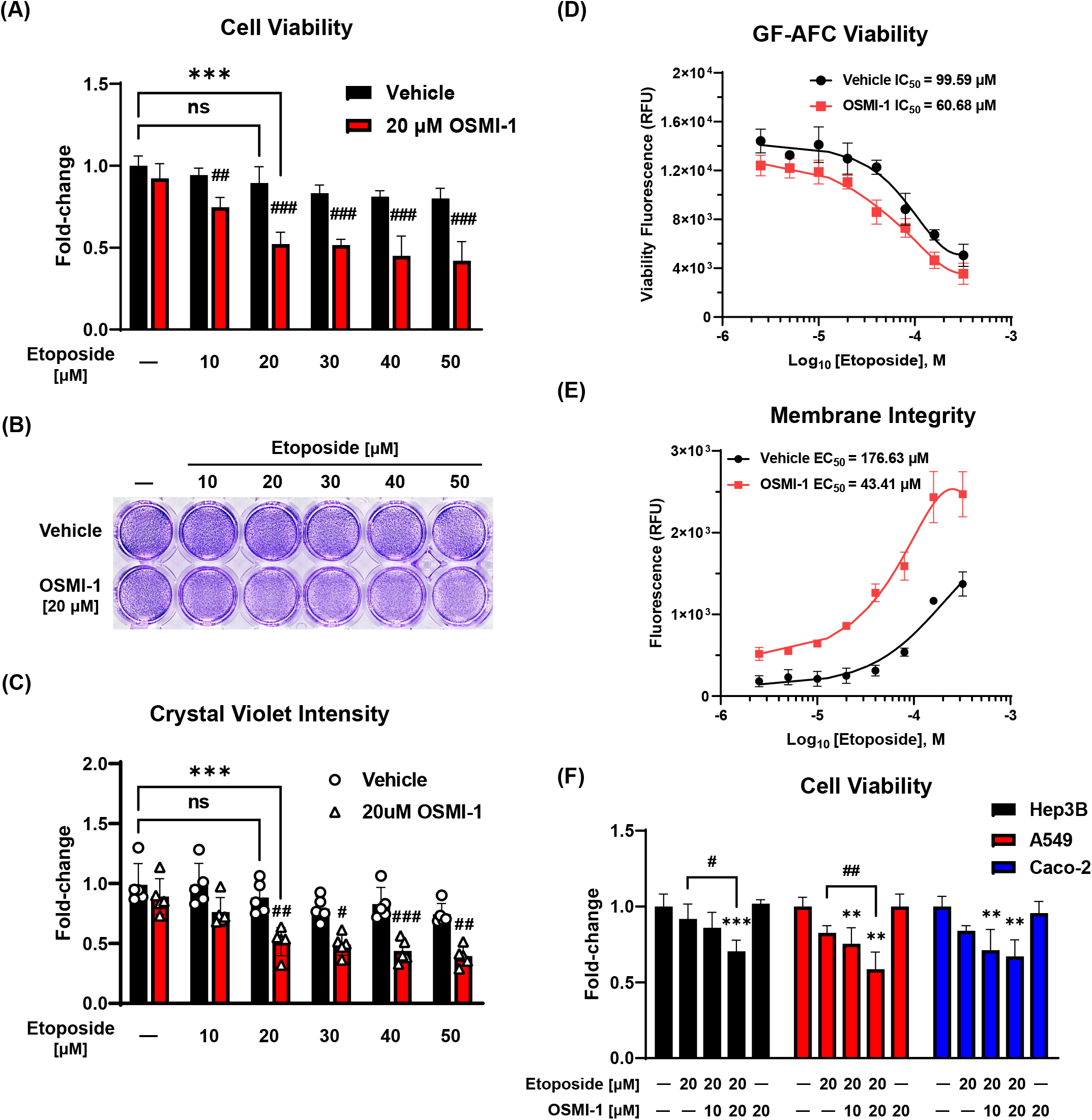
OSMI-1 enhances the sensitivity of HepG2 cells to etoposide. (A-C) Effect of etoposide combined with OSMI-1 on HepG2 human liver cancer cell viability assessed through the CCK viability assay (*n* = 5) and crystal violet staining (*n* = 5) 24 h post-treatment. Cotreatment with 20 μM OSMI-1 significantly reduced cell viability at all etoposide concentrations compared with etoposide alone. *** *P* < 0.001 versus the control group; # *P* < 0.05, ## *P* < 0.01, and ### *P* < 0.001 versus the vehicle control in the etoposide treatment group at the same concentration (mean ± SD). (D) Viability analysis via GF-AFC revealed a 1.64-fold reduction in the IC_50_ value of etoposide, from 99.59 to 60.68 μM, with 20 μM OSMI-1 (*n* = 3). (E) Membrane integrity assays showed a 4.07-fold decrease in the EC_50_ value of etoposide, from 176.63 to 43.41 μM, when combined with 20 μM OSMI-1 (*n* = 3). (F) Cell viability assays in Hep3B human liver hepatocellular carcinoma, A549 human lung carcinoma, and Caco-2 human colorectal adenocarcinoma cell lines confirmed a greater reduction in viability with the etoposide and OSMI-1 cotreatment (*n* = 4). ** *P* < 0.01 and *** *P* < 0.001 versus the control group; # *P* < 0.05 and ## *P* < 0.01 versus etoposide treatment alone (mean ± SD). Data are means from at least three independent experiments. Significance statistical analysis was performed with one-way or two-way ANOVA with *post hoc* Tukey test. ns, not significant.

### Etoposide decreases O-GlcNAcylation without directly interacting with OGT

To investigate the effect of etoposide on O-GlcNAc regulation, HepG2 cells were treated with etoposide for 24 h, and global O-GlcNAc expression was analyzed via western blot and immunocytochemistry analyses. Results revealed that etoposide downregulated O-GlcNAc expression in a concentration-dependent manner (Fig. 2A and B). When etoposide and OSMI-1 were administered either alone or in combination for 24 h, both treatments reduced O-GlcNAc expression, but cotreatment only reduced it to the same extent as OSMI-1 alone (Fig. 2C). To determine whether these O-GlcNAc modulatory effects were due to direct interaction with the OGT enzyme, an OGT inhibitor assay was conducted. The findings showed that etoposide did not significantly reduce OGT activity at concentrations below approximately 200 μM, whereas OSMI-1 had an IC_50_ value of 4.47 μM (Fig. 2D and E).

**Fig. 2.**
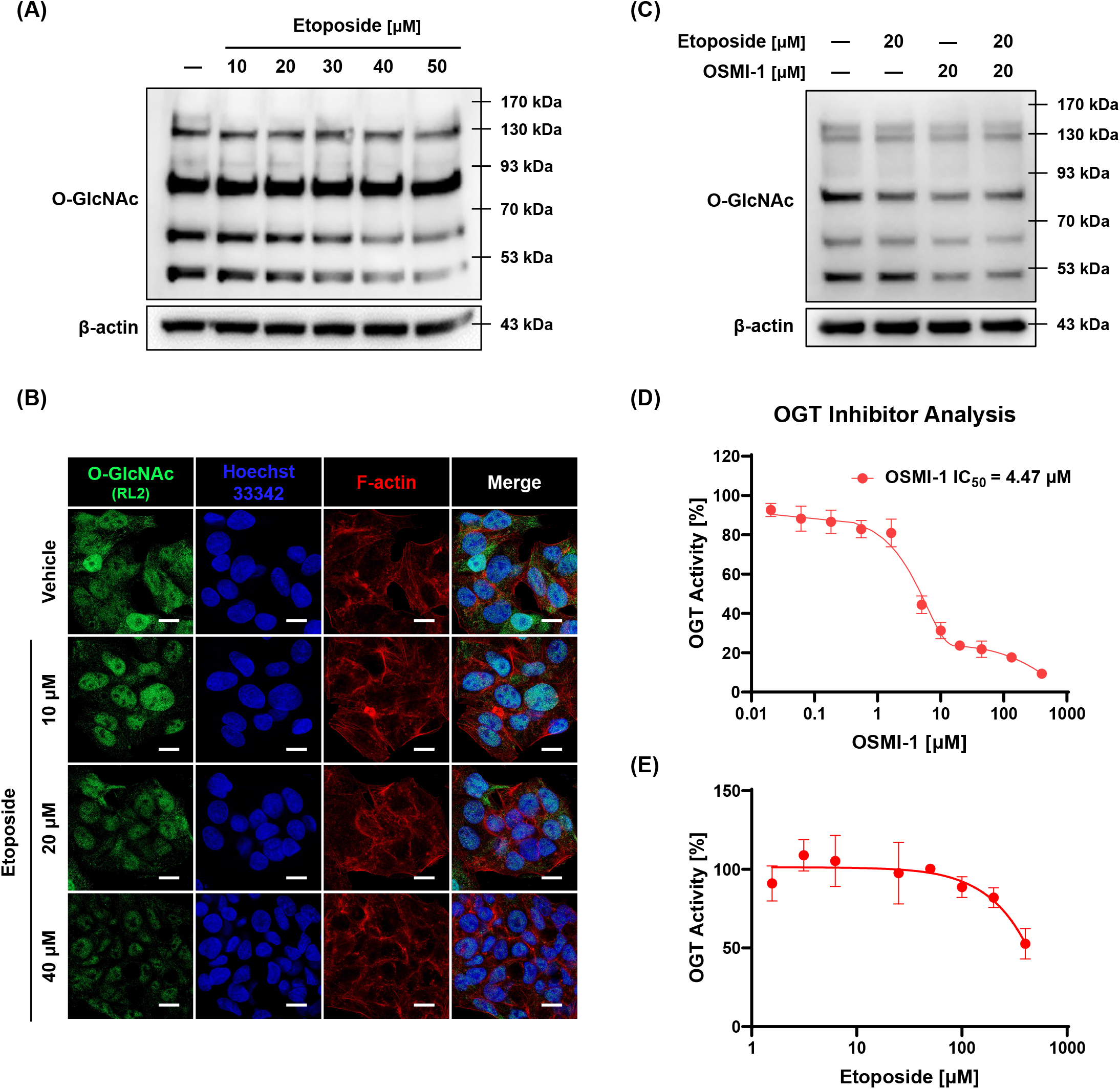
Etoposide reduces O-GlcNAcylation in a dose-dependent manner without directly interacting with OGT. (A) Etoposide treatment for 24 h resulted in dose-dependent reduction in O-GlcNAc expression in HepG2 cells. (B) Immunocytochemistry with O-GlcNAc antibody (RL2) (white scale bar: 10 μm) confirmed the findings in (A). (C) Etoposide and OSMI-1 cotreatment reduced O-GlcNAc expression levels comparable to OSMI-1 treatment alone. (D, E) OGT inhibition assays showed that etoposide did not significantly reduce OGT activity at concentrations below ~200 μM, whereas OSMI-1 exhibited an IC_50_ of 4.47 μM (*n* = 3; mean ± SD). Thus, etoposide decreases O-GlcNAc dose-dependently without directly interacting with OGT. Data are means from at least three independent experiments.

### Anticancer effect of etoposide occurs through downregulation O-GlcNAc expression

To assess whether the anticancer effect of etoposide is regulated by global changes in O-GlcNAc expression, we evaluated HepG2 cell viability using the OGT inhibitor OSMI-1 and the OGA inhibitor Thiamet G. Results indicated that OSMI-1 significantly enhanced the anticancer effect of etoposide in a dose-dependent manner, whereas Thiamet G did not induce significant changes (Fig. 3A). These findings were confirmed through crystal violet staining (Fig. 3B). Although the etoposide-induced decrease in crystal violet intensity was partially reversed by Thiamet G treatment, this increase did not achieve statistical significance (Fig. 3C). Further investigation of O-GlcNAc expression via western blot and immunocytochemistry analyses after 24 h of treatment revealed that O-GlcNAc expression was more prominently reduced when cells were treated with OSMI-1 compared to etoposide alone, and Thiamet G restored the etoposide-induced reduction in O-GlcNAc expression (Fig. 3D and E). Additionally, etoposide alone or in combination with OSMI-1 reduced OGA expression in HepG2 cells, and Thiamet G restored the OGA expression lowered by etoposide. However, OGT expression remained unaffected. These results suggest that the potentiation of etoposide’s anticancer effects by OSMI-1 occurs through O-GlcNAc downregulation.

**Fig. 3.**
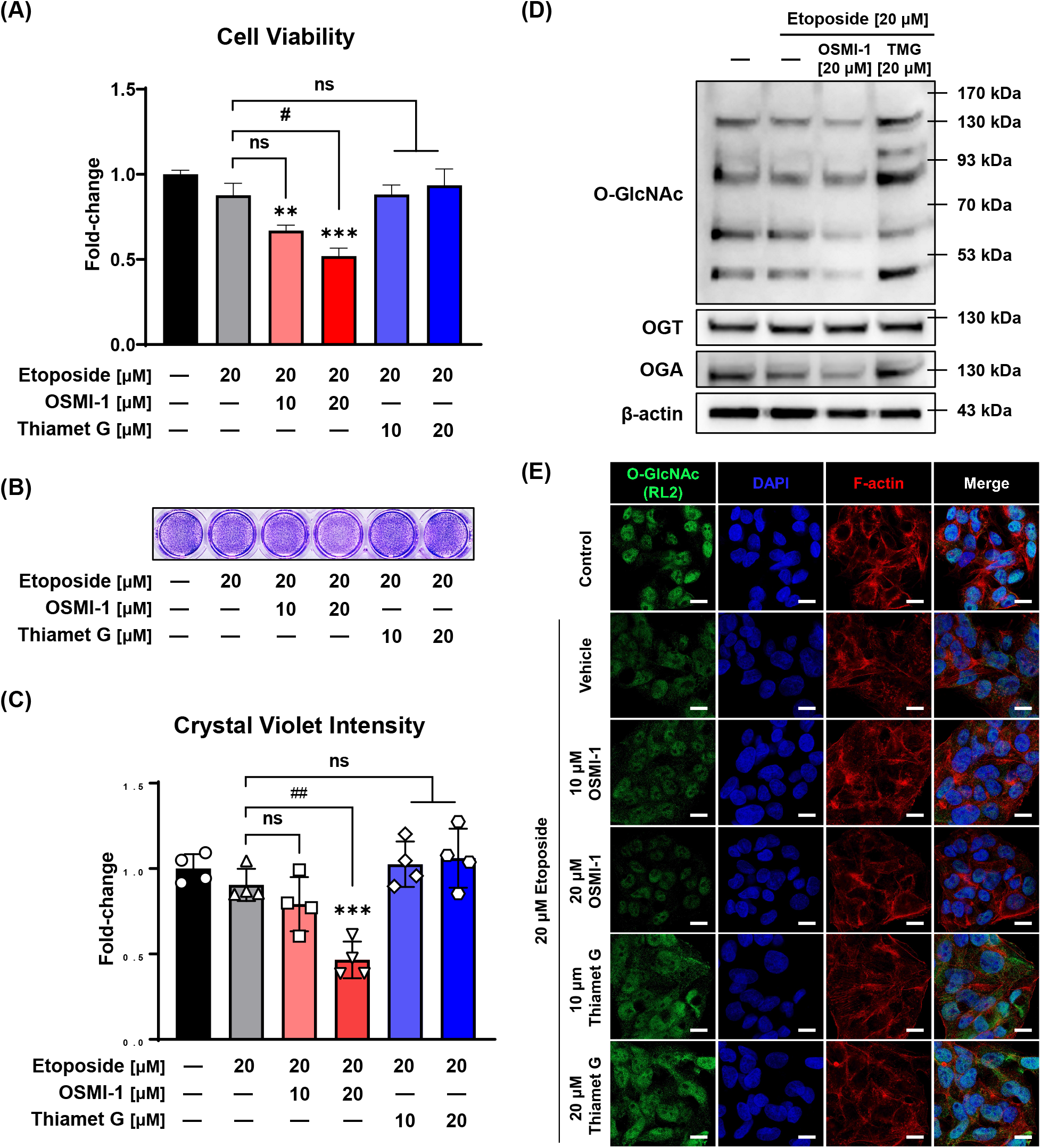
Etoposide’s anticancer effects are mediated by downregulation of O-GlcNAc expression. (A) Cell viability assays demonstrated that OSMI-1 (an OGT inhibitor) significantly increased the anticancer effect of etoposide in a concentration-dependent manner, whereas Thiamet G (an OGA inhibitor) did not alter viability (*n* = 4; mean ± SD). (B, C) Crystal violet staining assay corroborated the findings in (A) (*n* = 4; mean ± SD). (D, E) Western blot and immunocytochemistry analyses at 24 h post-treatment revealed that etoposide and OSMI-1 cotreatment reduced O-GlcNAc expression levels more than etoposide treatment alone. Thiamet G restored the O-GlcNAc expression levels lowered by etoposide (white scale bar: 10 μm). ** *P* < 0.01 and *** *P* < 0.001 versus the control group; # *P* < 0.05 and ## *P* < 0.01 versus the etoposide treatment alone. Data are the means from at least three independent experiments. Significance statistical analysis was performed with one-way ANOVA with *post hoc* Tukey test. ns, not significant.

### OGT inhibitor significantly increases etoposide-induced apoptosis

To determine if the enhancement of etoposide’s anticancer effect by OSMI-1 mediates the apoptosis pathway, fluorescence-activated cell sorting (FACS) analysis was performed. Annexin V was used to detect apoptotic cells, and the results revealed that cotreatment with 10 or 20 μM OSMI-1 and 20 μM etoposide for 24 h increased the proportion of cells in early apoptosis by 1.42- and 1.62-fold, respectively, compared with etoposide alone (Fig. 4A). However, 7-AAD staining did not significantly increase following 24 h of treatment. A luminescence assay was conducted to measure Annexin V binding to phosphatidylserine, yielding an EC_50_ value of 27.52 μM for the etoposide + 20 μM OSMI-1 cotreatment, which was 2.25-fold lower than the 61.94 μM value for etoposide treatment alone (Fig. 4B). Notably, etoposide and OSMI-1 cotreatment induced significant phosphatidylserine translocation from 4 h onward, with significant apoptosis occurring from 12 h onward, compared with etoposide treatment alone (Fig. 4C). Furthermore, TUNEL assay results confirmed that OSMI-1 treatment increased etoposide-induced apoptosis (Fig. 5A and B).

**Fig. 4.**
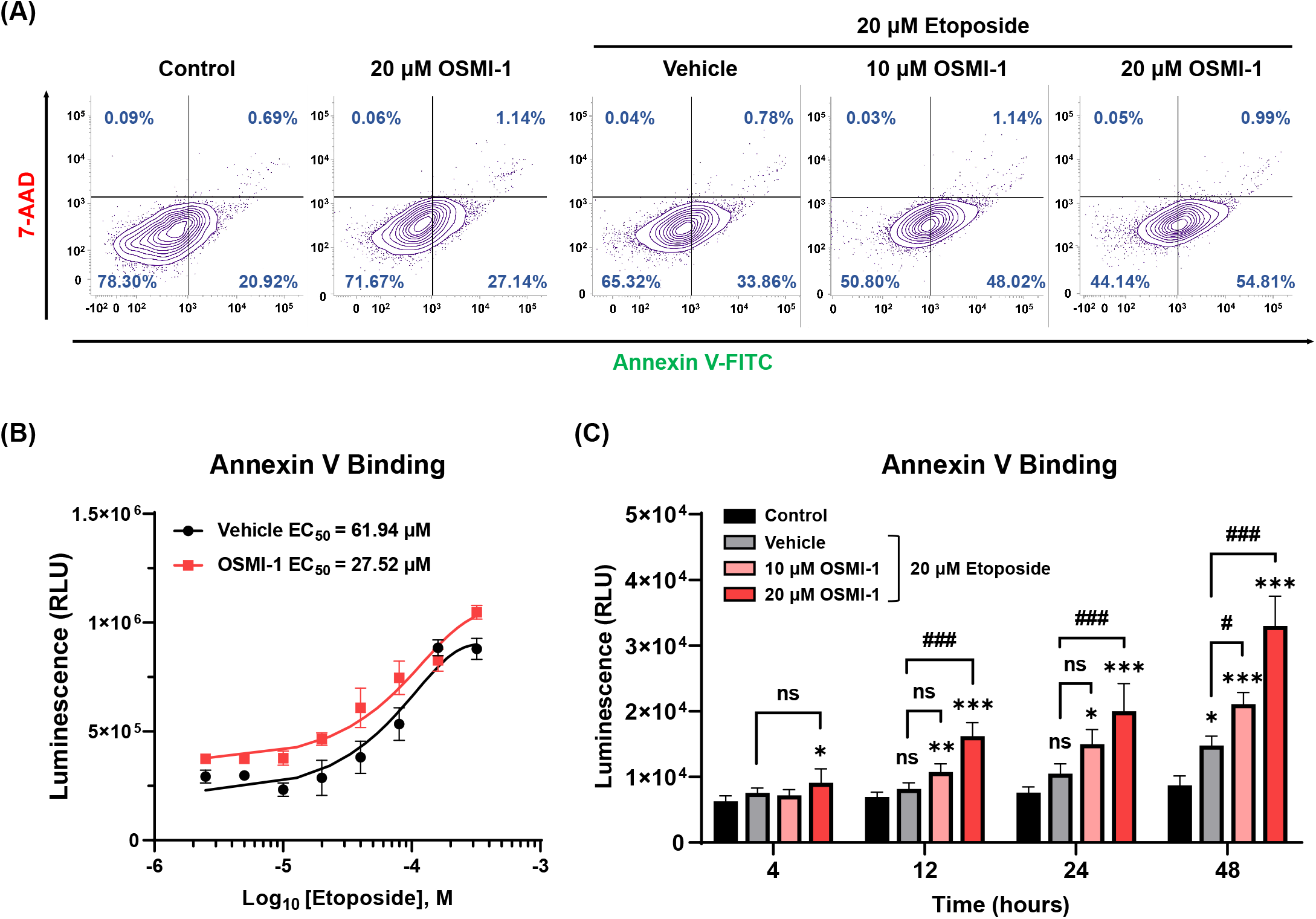
OSMI-1 significantly enhances etoposide-induced apoptosis. (A) FACS analysis using Annexin V and 7-AAD staining revealed that cotreatment with OSMI-1 and etoposide for 24 h significantly increased early apoptosis compared with etoposide treatment alone. (B) Quantitative luminescence analysis showed a 2.25-fold reduction in the EC_50_ of etoposide, from 61.94 to 27.52 μM, with 20 μM OSMI-1 (*n* = 3). (C) The apoptosis-inducing effect of the cotreatment was significantly greater than that of the control group from 4 h and that of etoposide treatment alone from 12 h post-treatment (*n* = 4). * *P* < 0.05, ** *P* < 0.01, and *** *P* < 0.001 versus the control group at each time point; ### *P* < 0.001 versus the etoposide treatment alone at each time point (mean ± SD). Data are means from at least three independent experiments. Significance statistical analysis was performed with one-way ANOVA with *post hoc* Tukey test. ns, not significant.

**Fig. 5.**
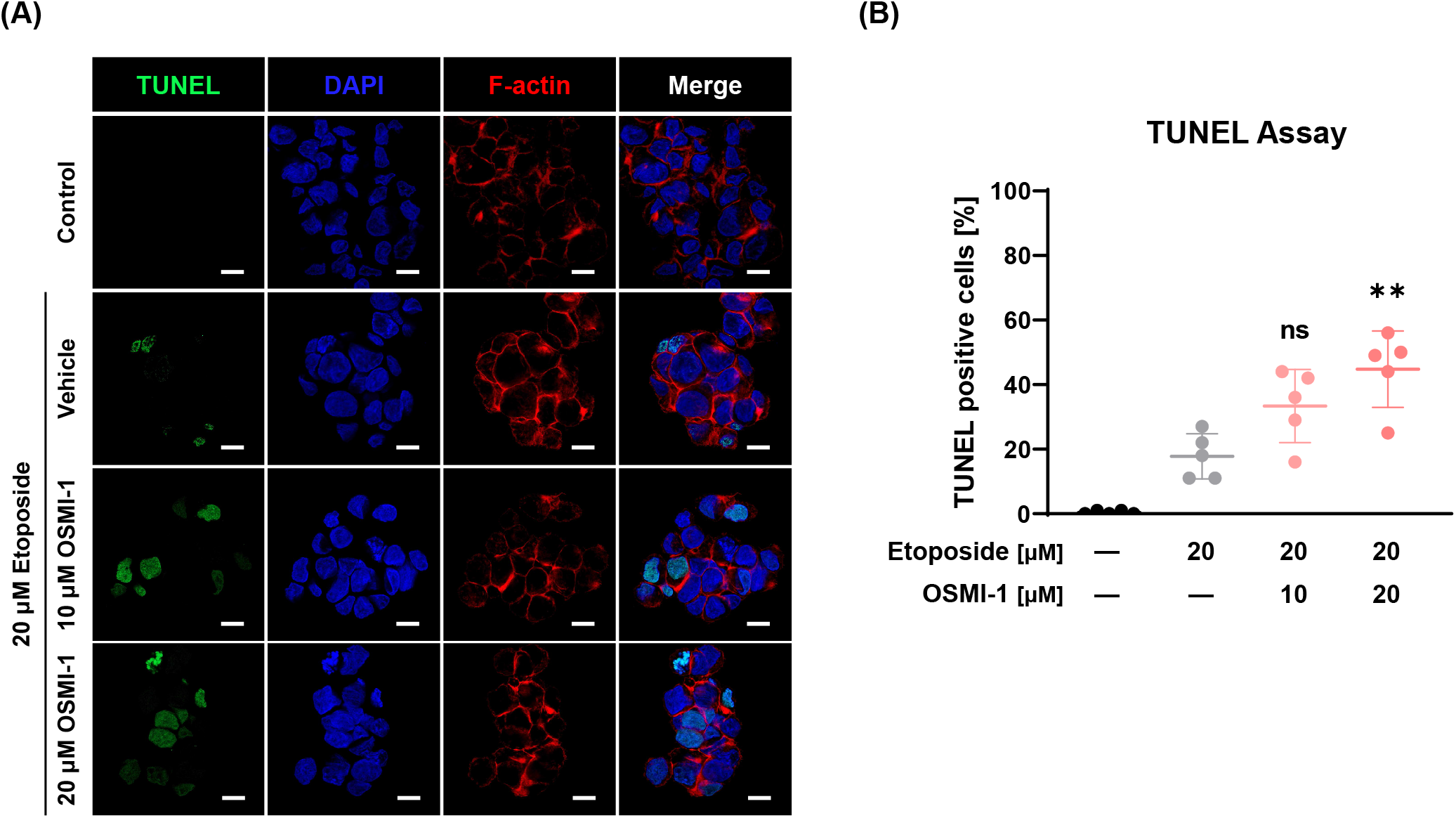
Etoposide and OSMI-1 cotreatment induces DNA fragmentation during apoptosis. (A, B) TUNEL assays demonstrated dose-dependent DNA fragmentation in HepG2 cells treated with etoposide and OSMI-1 for 24 h. OSMI-1 induced double DNA breaks in a dose-dependent manner. Thus, reduced O-GlcNAcylation amplifies etoposide-mediated apoptosis induction (*n* = 5; white scale bar: 10 μm). ** *P* < 0.01 versus the etoposide treatment alone (mean ± SD). Significance statistical analysis was performed with one-way ANOVA with *post hoc* Tukey test. ns, not significant.

### Synergistic anticancer effect of the OGT inhibitor is achieved through enhancement of the p53-mediated apoptotic pathway

To elucidate the mechanism underlying the synergistic anticancer effects of etoposide and the OGT inhibitor OSMI-1 p53 and cleaved-PARP1 expression levels were evaluated via western blot analysis following 24 h of treatment. Results showed that cotreatment with etoposide and 20 μM OSMI-1 significantly increased the expression of p53 and cleaved-PARP1 compared to etoposide treatment alone (Fig. 6A). Additionally, a caspase-3/7 activity assay, quantified via luminescence, revealed that etoposide alone enhanced caspase-3/7 activity in a dose-dependent manner, this activity was significantly increased further under the etoposide + 20 μM OSMI-1 cotreatment (Fig. 6B). These findings suggest that the enhanced downregulation of O-GlcNAcylation achieved by OSMI-1 potentiates activation of the p53-mediated apoptotic pathway, thereby augmenting the anticancer effects of etoposide.

**Fig. 6.**
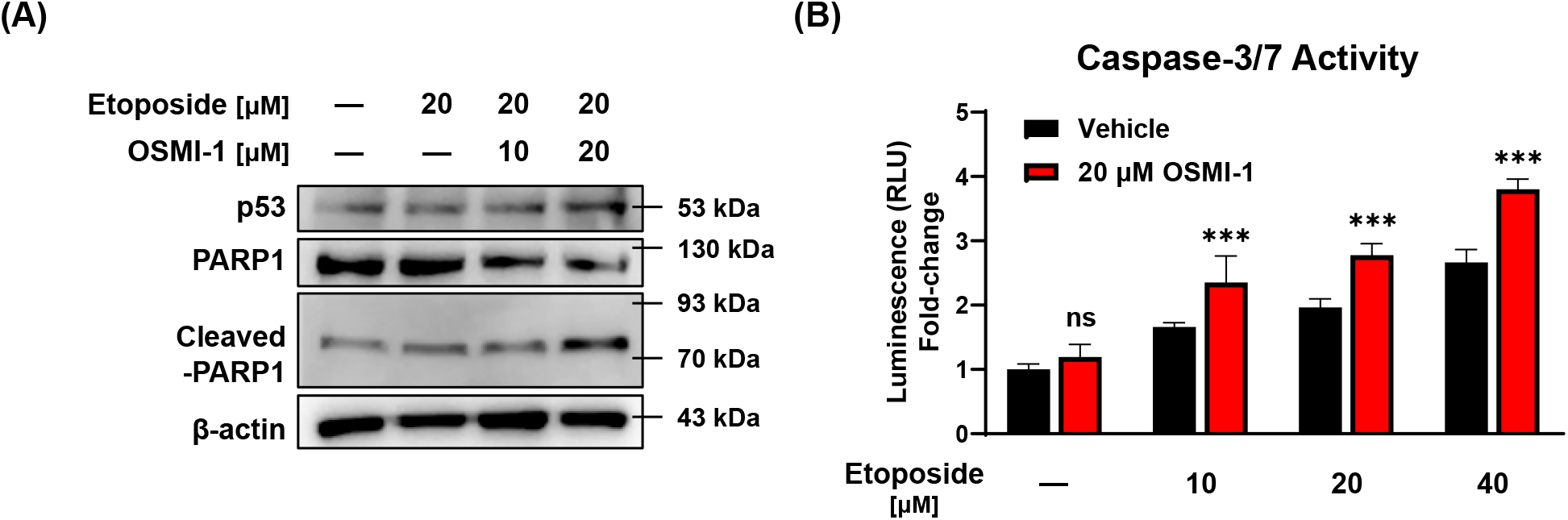
OSMI-1 enhances etoposide’s anticancer effect via activation of the p53/caspase-3/PARP1 pathway. (A) Western blot analysis revealed increased expression of p53 and cleaved-PARP1 (apoptosis markers) with 24 h of 20 μM etoposide and 20 μM OSMI-1 cotreatment compared to etoposide alone. (B) Quantitative luminescence analysis showed that caspase-3/7 activity was also higher with etoposide and 20 μM OSMI-1 cotreatment (*n* = 4). *** *P* < 0.001 versus the etoposide treatment alone (mean ± SD). Data are means from at least three independent experiments. Significance statistical analysis was performed with two-way ANOVA with *post hoc* Tukey test. ns, not significant.

## Discussion

Etoposide, a plant-derived alkaloid, inhibits cell proliferation across various adenocarcinoma cell types by arresting cell division. Mammalian cells express two Topo II isoenzymes, namely Topo IIα and β, with Topo IIα often overexpressed in tumors, serving as a cell proliferation marker [28]. The primary cytotoxicity mechanism of etoposide involves targeting Topo IIα, which underpins its selective action on cancer cells [29]. However, tumors resistant to Topo II inhibitors and cell lines with decreased Topo II expression exhibit reduced sensitivity to etoposide, suggesting a direct correlation between resistance and reduced Topo IIα expression [30]. Consequently, combining Topo II inhibitors, including etoposide, with agents acting via alternative mechanisms of action has become a common strategy to counteract resistance.

Our findings demonstrate that OSMI-1 significantly enhanced the sensitivity of HepG2 cells to etoposide (Fig. 1D and E). Notably, the EC_50_ value for membrane integrity, a marker of cytotoxicity, decreased by more than 4-fold with OSMI-1 cotreatment. This result was validated by a marked reduction in cell viability under 10 μM etoposide treatment combined with OSMI-1 compared to etoposide alone (Fig. 1A). Although additional research is required to clarify the precise effect of reduced O-GlcNAcylation on etoposide’s interaction with Topo II or subsequent molecular cascades, our study suggests that inhibiting O-GlcNAcylation may serve as an adjuvant strategy to enhance etoposide’s anticancer efficacy. Previous studies have shown that reducing excessive O-GlcNAcylation can enhance the effectiveness of anticancer drugs [25, 26]. For example, the O-GlcNAcylation of the Topo IIα Ser1469 residue in breast cancer cells has been implicated in increased drug resistance to the chemotherapeutic drug adriamycin, and its inhibition has been shown to enhance therapeutic outcomes *in vitro* and *in vivo* [31]. This is consistent with findings that cancer cell lines and human tumors frequently exhibit elevated O-GlcNAcylation [32, 33]. In line with previous literature, our findings demonstrated that elevating O-GlcNAcylation via Thiamet G tended to decline the anticancer effect of etoposide (Fig. 3A and C). These observations suggest a potential causal relationship between heightened O-GlcNAcylation and tumorigenesis, although an analysis of 33 cancer types in the Cancer Genome Atlas database showed that mutations in O-GlcNAcylation cycling enzymes are rare compared with highly mutated genes, such as *p53* and *PIK3CA* [34], indicating that increased O-GlcNAcylation may be a phenotypic consequence of cancer progression.

Limited evidence exists on the effects of Topo II inhibitors on O-GlcNAcylation regulation. Doxorubicin, another Topo II inhibitor, has been reported to reduce O-GlcNAcylation in Cos-7 cells [35], but such effects have not been studied extensively for etoposide. Cotreatment with doxorubicin and OSMI-1 was previously shown to induce excessive apoptosis via the p53/PARP1 pathway in HepG2 cells [25], similar to our findings with etoposide in this study. However, studies using *Xenopus* egg extracts revealed that etoposide and doxorubicin exhibit differing Topo II inhibition mechanisms: etoposide traps Topo II behind replication forks in a Topo II–dependent manner, whereas doxorubicin stops the replication forks by inserting directly into the parental DNA in a Topo II–independent manner [36]. Therefore, the present study provides new insights into the interplay between Topo II inhibitors and O-GlcNAcylation regulation.

Our results confirmed that etoposide treatment reduces O-GlcNAcylation in a dose-dependent manner without directly interacting with OGT (Fig. 2A, C, and (E). Although high etoposide concentrations (400 μM) appeared to reduce OGT activity by approximately 47.3%, this inhibitory effect was likely exaggerated owing to etoposide solubility issues (Fig. 2E). These findings suggest that the apoptotic cascade initiated by etoposide’s inhibition of Topo II is associated with O-GlcNAcylation downregulation. This hypothesis is supported by the dose-dependent increase in apoptosis observed with OSMI-1 cotreatment (Fig. 4A and (5B). Interestingly, although etoposide treatment alone did not significantly alter OGT expression in HepG2 cells, it reduced OGA levels, an effect further amplified by OSMI-1 cotreatment (Fig. 3D). This may reflect a compensatory mechanism aimed at restoring reduced O-GlcNAcylation levels, as reported in a prior study [37].

Under cellular stress, whether arising internally or from external factors, p53 expression and stability are elevated, triggering the activation of genes responsible for cell cycle arrest, DNA repair, apoptosis, and autophagy [38]. Following DNA damage, the cell’s fate depends on its severity, with damage either repaired or cell death (e.g., apoptosis and necrosis) being initiated. In cases of minimal damage, the cell cycle is arrested, enabling the activation of DNA repair mechanisms. PARP1 plays a crucial role in genome integrity by recognizing DNA damage and facilitating the repair pathway [39]. However, when DNA damage becomes irreparable, PARP1 is cleaved by caspase-3/7, a process regulated by the p53 pathway. This cleavage inactivates PARP1, leading the cell toward apoptosis. [40]. Our findings demonstrated that cotreatment with OSMI-1 enhanced activation of the p53/PARP1 pathway to a greater extent than etoposide treatment alone (Fig. 6A). This molecular insight aligns with the observed leftward shift in the phosphatidylserine–Annexin V binding curve under etoposide + 20 μM OSMI-1 cotreatment conditions (Fig. 4B). Furthermore, the synergistic effect of OSMI-1 was amplified over time compared with etoposide alone (Fig. 5C).

Etoposide has been extensively used in drug screening studies owing to its diverse cellular effects, which include induction of autophagy, cell senescence, and apoptosis [41, 42]. These properties highlight the need for further studies on the combination effects and mechanisms of etoposide and OSMI-1 across broader indications. As OSMI-1 functions as an OGT inhibitor, downregulating global O-GlcNAcylation, future studies should focus on identifying the specific O-GlcNAcylated residues in apoptosis-related proteins to clarify the molecular mechanisms underlying this synergy. Although this study primarily used HepG2 human liver hepatocellular carcinoma, the fundamental mechanism of etoposide’s action is consistent across various cell types. This suggests that similar effects can be anticipated in other cancers. Consistent with this hypothesis, we confirmed the enhanced anticancer effect of the etoposide–OSMI-1 combination in Hep3B human liver hepatocellular carcinoma, A549 human lung carcinoma, and Caco-2 human colorectal adenocarcinoma cell lines (Fig. 1F). It should be confirmed whether these effects can be maintained *in vivo*, since our study did not verify its efficacy in an animal model. Etoposide is a chemotherapeutic option used primarily for the treatment of lung or testicular cancer; therefore, *in vivo* study using these other tumors could be considered based on existing studies. Additionally, animal studies should be conducted to adjust the dosing ratio of etoposide and OSMI-1 to maximize efficacy. Given that OGT is an important enzyme in systemic homeostasis, cotreatment of OSMI-1 may result in adverse effects. Therefore, future *in vivo* studies should be designed to simultaneously verify safety and anticancer effects.

In conclusion, downregulation of global O-GlcNAcylation is integral to the apoptotic induction process initiated by etoposide. Additional downregulation of O-GlcNAcylation via OGT inhibition significantly amplifies etoposide’s anticancer effects through activation of the p53-mediated pathway (Fig. 7). Although further *in vivo* studies and molecular analyses are necessary to validate and elucidate this synergistic effect and its mechanism, our results suggest that targeting O-GlcNAcylation presents a promising approach for combination chemotherapy in cancer treatment.

**Fig. 7.**
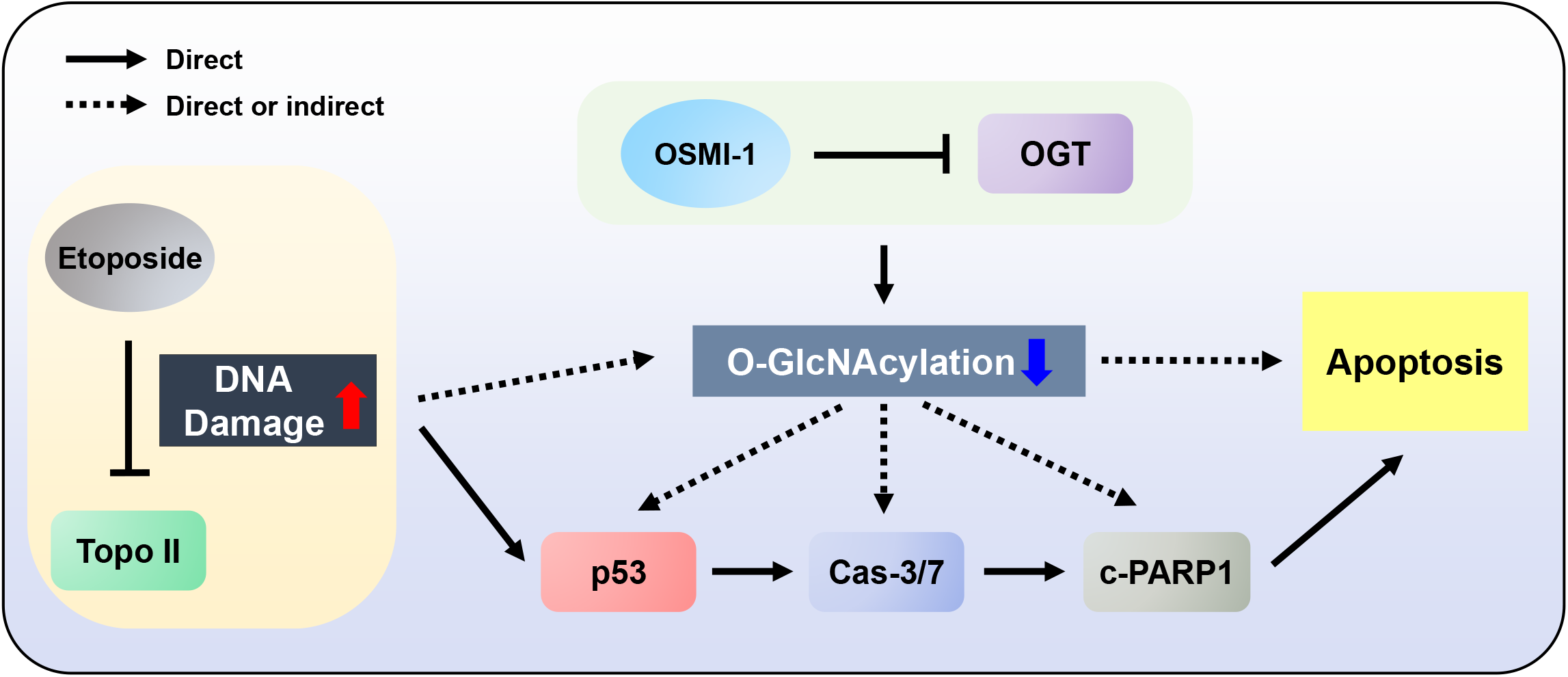
Downregulation of O-GlcNAcylation augments etoposide-induced p53-mediated apoptosis. Etoposide induces apoptosis by forming a ternary complex with topoisomerase II (Topo II) and DNA, inhibiting DNA strand religation and causing DNA damage. This activates the p53-mediated apoptosis pathway, increasing caspase-3/7 (Cas-3/7) activity. Activated caspase-3/7 cleaves PARP1, leading to apoptosis and thereby maintaining genome integrity. Although etoposide downregulates O-GlcNAcylation during this process, the exact mechanism remains unclear. Cotreatment with OSMI-1, an OGT inhibitor, further suppresses O-GlcNAcylation, enhancing etoposide’s p53-mediated apoptosis pathway.

## Abbreviations

ANOVA: analysis of variance
ATCC: American Type Culture Collection)
BSA: bovine serum albumin
Cas-3/7: caspase-3/7
EMEM: Eagle’s Minimum Essential Medium
OGA: O-GlcNAcase
O-GlcNAc: O-linked β-N-acetylglucosamine
O-GlcNAcylation: O-linked β-N-acetylglucosamine modification
OGT: O-GlcNAc transferase
PBS: phosphate-buffered saline
PBST: PBS containing 0.1% Tween 20
P/S: penicillin/streptomycin
SD: standard deviation
TBST: 0.05% Tween 20 Tris-buffered saline
Topo: II topoisomerase II
UDP: uridine diphosphate

## Data accessibility

The data that support the findings of this study are available from the corresponding author upon reasonable request.

## Author contributions

JL and GK designed the experiments under the supervision of BH and SL. JL wrote the manuscript. JL, JP, and MK performed the experiments and analyzed the data. JL, GK, and BH performed literature research. JL designed the research template. All authors have read and agreed to the published version of the manuscript.

## Acknowledgements

This study was funded by the R&D Headquarter of Korea Ginseng Corporation

